# Structure-Guided Mutagenesis Targeting Interactions between pp150 Tegument Protein and Small Capsid Protein Identify Five Lethal and Two Live Attenuated HCMV Mutants

**DOI:** 10.1101/2024.01.22.576707

**Authors:** Alex Stevens, Ruth Cruz-cosme, Najealicka Armstrong, Qiyi Tang, Z. Hong Zhou

**Affiliations:** California NanoSystems Institute, University of California, Los Angeles, CA 90095, USA; Department of Chemistry and Biochemistry, University of California, Los Angeles, CA 90095, USA; Department of Microbiology, Immunology and Molecular Genetics, University of California, Los Angeles, CA 90095, USA; Department of Microbiology, Howard University College of Medicine, Washington, DC 20059, USA

## Abstract

Human cytomegalovirus (HCMV) replication relies on a nucleocapsid coat of the 150kDa, subfamily-specific tegument phosphoprotein (pp150) to regulate cytoplasmic virion maturation. While recent structural studies revealed pp150-capsid interactions, the role of specific amino-acids involved in these interactions have not been established experimentally. In this study, pp150 and the small capsid protein (SCP), one of pp150’s binding partners found atop the major capsid protein (MCP), were subjected to mutational and structural analyses. Mutations to clusters of polar or hydrophobic residues along the pp150-SCP interface abolished viral replication, with no replication detected in mutant virus-infected cells. Notably, a single point mutation at the pp150-MCP interface significantly attenuated viral replication, unlike the situation of pp150-deletion mutation where capsids degraded outside host nuclei. These functionally significant mutations targeting pp150-capsid interactions, particularly the pp150 K255E replication-attenuated mutant, can be explored to overcome the historical challenges of developing effective antivirals and vaccines against HCMV infection.

## 1. Introduction

Human cytomegalovirus (HCMV), is a ubiquitous beta-herpesvirus which exhibits typical seroprevalence of ∼83% (Zuhair et al., 2019). Though innocuous in healthy adults, HCMV infection is a leading causative agent of debilitating illnesses, such as retinitis or hepatitis, amongst the immunocompromised and the immunologically naïve (Manicklal et al., 2013). HCMV is also the leading viral cause of birth defects world-wide, often causing neurological disabilities such as deafness, blindness, and microcephaly (Manicklal et al., 2013). The prevalence of HCMV and the public health burden it represents, motivates ongoing work to develop more effective treatments by making use of newly available structural data.

HCMV is a member of the β-herpesvirus subfamily of the *Herpesviridae*, and shares general morphological characteristics of viruses from the α-herpesvirus (e.g., herpes simplex virus or HSV) and γ-herpesvirus (e.g., Kaposi’s sarcoma-associated herpesvirus or KSHV) subfamilies. The virion has four layers: the large double-stranded DNA genome is packed tightly into the icosahedral capsid, which is coated in amorphous tegument proteins and enclosed within a lipid envelope. The HCMV capsid is composed of 955 copies of the major capsid protein (MCP) arranged into 150 hexons with 11 pentons at the icosahedral 5-fold (I5) vertices. The 12^th^ I5 vertex is occupied by a unique portal vertex, through which the viral genome is translocated. 360 triplex protein complexes (Tri), each composed of one Tri1 and two Tri2 (Tri2A & Tri2B) copies, decorate the exterior of the MCP shell, riveting neighboring MCPs together, while small capsid protein (SCP) molecules tip each MCP tower (Figure 1A). SCP, which varies from 8kDa in HCMV and 15 kDa in KSHV, has the greatest structural and sequence variability amongst herpesviruses (Close et al., 2018; Dai et al., 2015), being dispensable among both α- and γ-herpesviruses but not β-herpesviruses (Borst et al., 2022; Dai et al., 2013).

**Figure 1.**
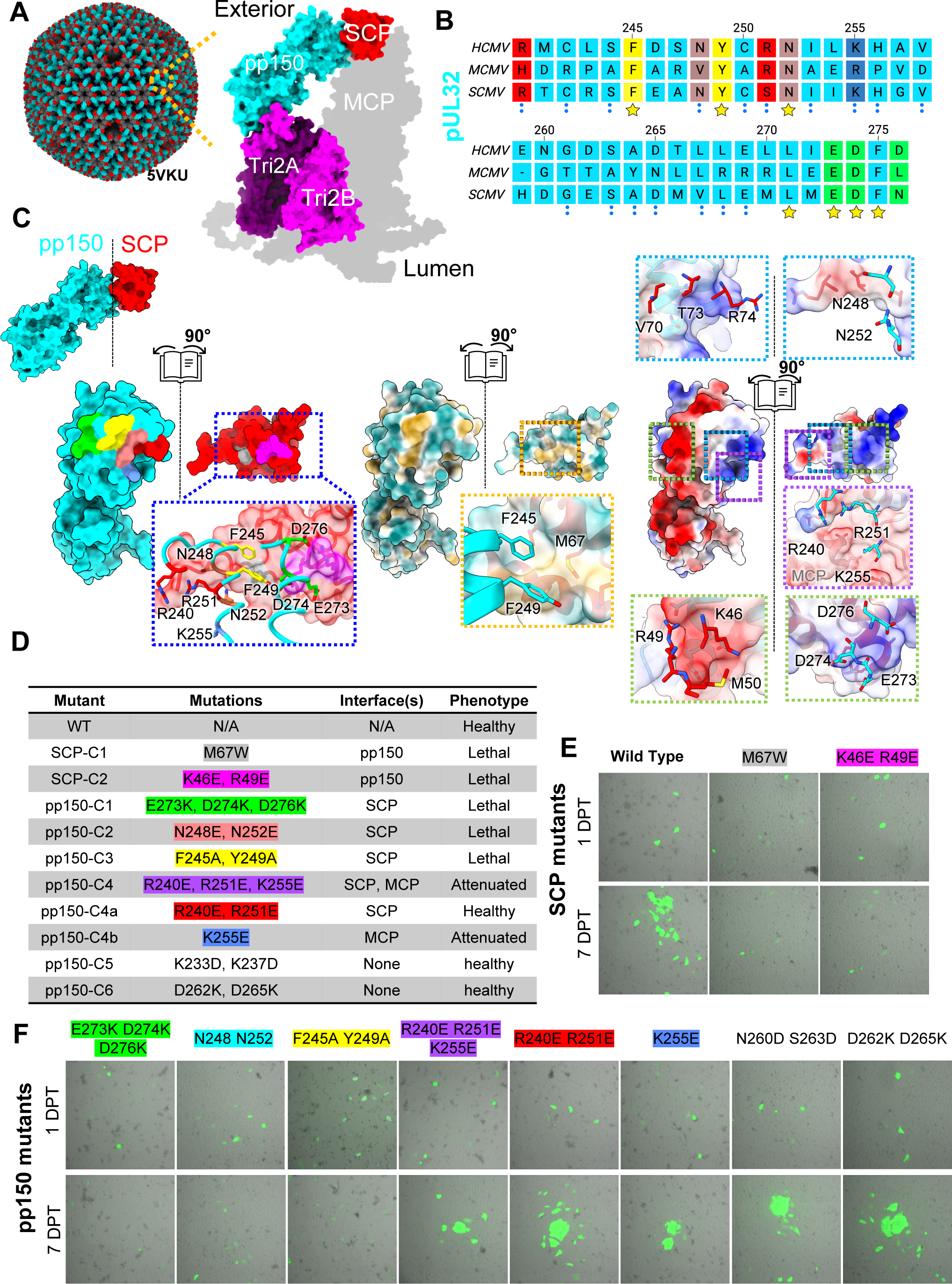
Structure-guided mutational analysis of the pp150-SCP interface. (A) Atomic model of the HCMV capsid with pp150 coat (left). The inset (right) shows zoomed-in view of subunits colored to emphasize different subunit interactions. (B) Sequence alignment of HCMV, murine CMV (MCMV), and simian CMV (SCMV) pp150 homologues between pp150 residues 240-276. (C) surface interfaces between pp150 and SCP, with mutations colored by clusters to enhance detail (left). Surfaces with Hydrophilic, neutral, and hydrophobic areas colored in cyan white and gold respectively (middle) while positive, neutral, and negative electrostatic potentials are shown in blue, white, and red respectively (right). (D) Table listing cluster mutations, and their effect on virus replication, as observed in transfected cells. (E) Representative Fluorescence microscopy images of cells transfected with SCP mutant BACs, 1 (top) or 7 (bottom) days post transfection (DPT). (F) Same as (E) but from pp150 mutant BACs at 1 or 7 DPT.

As a β-herpesvirus, HCMV has several features which differentiate it from members of the other two Herpesviridae subfamilies. For instance, with greater genomic density than its α-herpesvirus cousins (Bauer et al., 2013; Muller et al., 2021), the HCMV capsid is subject to internal pressures beyond the 18 atmospheres experienced by herpes simplex virus (HSV) (Brandariz-Nuñez et al., 2019). Strangely, in HCMV the stabilizing capsid vertex-specific components (CVSCs) present in HSV are replaced with the 150 kDa β-herpesvirus specific tegument phosphoprotein (pp150) (Dai and Zhou, 2018; Wang et al., 2018; Yu et al., 2017), despite HCMV CVSCs remaining at the portal vertex (Li et al., 2021; Liu et al., 2019; Sae-Ueng et al., 2014). Indeed, structural studies by our lab and others have found pp150 coats the capsid exterior of mature HCMV virions, bridging the gap between triplex heterotrimers and the small capsid protein (SCP) seated atop the neighboring major capsid proteins (MCP) of pentons and hexons (Fig. 1A) (Dai et al., 2013; Li et al., 2021; Liu et al., 2019).

Pp150 is thought to bind to nascent HCMV nucleocapsids in the nucleus and co-egress to the cytoplasm where it enhances capsid stability and regulates maturation events at the viral assembly complex (vAC) (AuCoin et al., 2006; Tandon and Mocarski, 2008; Tandon and Mocarski, 2011). The capsid bound, N-terminal region of pp150 consists of an upper and lower helix bundle with the later standing off Tri and the former contacting a neighboring SCP (Figure 1A) (Baxter and Gibson, 2001; Dai et al., 2013; Yu et al., 2017). Near the SCP and Tri binding sites of the helix bundles are two highly conserved regions (designated as CR1 & CR2) (Baxter and Gibson, 2001; Yu et al., 2017). Peptide mimetics of CR2 were shown to inhibit capsid binding of pp150 though the precise mechanism of inhibition remains unclear (Mitra et al., 2021). While other investigations including those from our lab have observed SCP to mediate pp150-capsid association (Dai et al., 2013; Yu et al., 2017)the interface between SCP and pp150 has yet to be thoroughly characterized and it remains unclear which, if any, residues or domains are implicated in pp150-capsid binding.

In this study, we utilized bacterial artificial chromosomes (BACs) to introduce structure guided mutations at the pp150-SCP interface and observed the impact on viral protein interactions and replication. Biochemical analysis and fluorescence imaging suggest mutations along the pp150-SCP interface invariably produce lethal phenotypes in HCMV which can likely be attributed to disruption of pp150-SCP association. Meanwhile, we identified a mutation at the pp150-MCP interface which significantly attenuates but does not entirely abolish viral replication. Through ultra-structural analysis, we observed no qualitative effect on capsid assembly, genome retention, or cytoplasmic maturation events, suggesting that this mutation to pp150-MCP interface may impact replication following egress from the cell. We also provide the first structural evidence to support previous assertions that pp150 associates with capsids within the nucleus of infected cells and preferentially binds genome containing capsids.

## 2. Materials and Methods

### 2.1 Bacterial artificial chromosome mutagenesis of HCMV AD169WT

To mutate the pp150 and SCP, we employed a seamless BAC technique using galK as the selection marker (Warden et al., 2011). Briefly, the BACmid of wild type AD169, vDW215-BADrUL131 (Wang and Shenk, 2005) was transformed into E. coli SW102. A galK DNA fragment that was made from pgalK by PCR and contains ends homologous to the open reading frame (ORF) of pp150 or SCP was electroporated into SW102 (harboring vDW215-BADrUL131) to replace pp150 or SCP by homologous recombination, resulting in the following BACmid: vDW215-BADrUL131-pp150galK or vDW215-BADrUL131-SCPgalK. Then the galK DNA was replaced with a PCR fragment composed of pp150 or SCP DNA, from which fragment the indicated mutations were made, resulting in vDW215-BADrUL131-pp150 E273K D274K D276K, vDW215-BADrUL131-pp150 F245A Y249A, vDW215-BADrUL131-pp150 R240E R251E K255E, vDW215-BADrUL131-pp150-N248E N252E, vDW215-BADrUL131-pp150 K255E, vDW215-BADrUL131-pp150 R240E R251E, vDW215-BADrUL131-pp150 N260D S263D, vDW215-BADrUL131-pp150 D262K D265K, vDW215-BADrUL131-SCP K46E R49E, and vDW215-BADrUL131-SCP M67W. The resultant BACmids were transfected into ARPE-19 cells (ATCC® CRL-2302™) to make respective viruses. The BACmids and viruses were verified by restriction enzyme digestion, DNA sequencing, and PCR. The complete pp150 and SCP gene was sequenced and confirmed to be correct.

### 2.2 Molecular cloning

To express PP150 and SCP proteins, we performed cloning by inserting the pp150 or SCP gene DNA into the linearized pcDNA3 vector using BamHI and HindIII enzymes. DNA fragments were produced through PCR, utilizing BAC DNAs (wild type or mutant) as templates. During PCR, we incorporated an HA tag at the C-terminus of pp150 and a FLAG tag at the N-terminus of SCP. Subsequently, these DNA fragments were ligated with the linearized pcDNA3 vector using DNA ligase. Following transformation into E. coli, colonies were selected and verified via electrophoresis in an agarose gel. The wild type (WT) and mutant plasmids were confirmed through Sanger sequencing.

### 2.3 Co-Immunoprecipitation (Co-IP) assay and immunoblot analysis

Initially, PP150- and SCP-expressing plasmids were co-transfected into HEK293T cells using Lipofectamine 3000 Reagent (ThermoFisher Scientific, Cat# L3000015) and allowed to incubate for 24 hours. At 24 hours post transfection (hpt), cells were lysed. Lysis was performed using an ice-cold lysis buffer comprising 25 mM Tris-HCl (pH 7.4), 150 mM NaCl, 1% NP-40, 1 mM EDTA, and 5% glycerol. Additionally, this lysis buffer was supplemented with a protease inhibitor cocktail (Sigma, Cat# P8340), and the cell lysis process was performed on ice for a duration of 10 minutes. Following this, the lysates underwent centrifugation at 3,000 Xg for 5 minutes, and the resulting supernatants were meticulously transferred to new tubes.

For the subsequent immunoprecipitation step, these supernatants were subjected to overnight incubation at 4°C, in the presence of specific antibodies such as mouse anti-HA, and mouse anti-FLAG. These incubation mixtures were then combined with protein G-Sepharose beads (Amersham Pharmacia Biotech AB, Sweden), as per the manufacturer’s instructions, and allowed to further incubate for 3 hours.

Following immunoprecipitation, the beads were washed three times with PBS containing 0.1% bovine serum albumin, along with a protease inhibitor cocktail. The immune-precipitated complexes were then resuspended in a mixture of PBS and 2× Laemmli buffer (20 μl each). After a brief heating to 95°C for 5 minutes, the beads were separated by centrifugation, and the resultant supernatants were subjected to SDS-PAGE and subsequent immunoblotting.

For the immunoblot analysis, protein samples (ranging from 10 to 20 µg loaded in each lane) were separated utilizing a 7.5% polyacrylamide gel, followed by their transfer onto nitrocellulose membranes (Amersham Inc., Piscataway, NJ). Subsequently, these membranes were blocked with nonfat milk (5% in PBS pH 7.4) for a period of 60 minutes at room temperature. Thereafter, the membranes were incubated overnight at 4°C with primary antibodies. Post-primary antibody incubation, the membranes were treated with a horseradish peroxidase-coupled secondary antibody (Amersham Inc.). Protein detection was achieved through an enhanced chemiluminescence method (Pierce, Rockford, Ill.), following standard protocols. To facilitate the detection of additional proteins of interest, the membranes were stripped using a stripping buffer (comprising 100 mM β-mercaptoethanol, 2% SDS, and 62.5 mM Tris-HCl, pH 6.8), washed with PBS containing 0.1% Tween 20, and subsequently reused for subsequent different protein detection.

### 2.4 Immunofluorescence assay (IFA)

For IFA, ARPE-19 cells were cultured on coverslips and subsequently fixed with 1% paraformaldehyde for 10 minutes at room temperature. Following fixation, the cells were permeabilized by treatment with 0.2% Triton-X100 for 20 minutes on ice. Subsequently, a sequential series of incubations was performed. The cells were first incubated with primary antibodies, followed by incubation with Texas Red (TR)- or FITC-labeled secondary antibodies, all of which were dissolved in phosphate-buffered saline (PBS). Each incubation step lasted for 30 minutes. Finally, the cells were equilibrated in PBS, stained to visualize DNA using Hoechst 33258 at a concentration of 0.5 μg/ml, and then securely mounted using Fluoromount G (obtained from Fisher Scientific, Newark, Del.).

### 2.5 Confocal microscopy

We utilized a Leica TCS SPII confocal laser scanning system to examine the cells. During imaging, we recorded two or three channels either simultaneously or sequentially. We took special care to control for potential signal interference between the fluorescein isothiocyanate and Texas Red channels, as well as between the blue and red channels.

### 2.6 CryoEM of BAC derived nucleocapsids

HCMV nucleocapsids were prepared as described previously (Dai and Zhou, 2014). Briefly, ARPE-19 cells infected with HCMV-AD169+GFP BAC derived virus were seeded onto 18 x T175 flasks. When ∼80% of cells exhibited cytopathic effects of infection, or after ∼10 days in pp150-C4 mutants, cells were scrapped and pelleted. Pellets were washed and resuspended in PBS (pH 7.4) containing 1% NP40 solution and protease inhibitors (PI) 2 times to disrupt the cell membrane. Nuclei were pelleted and resuspended in 8 mL hypertonic lysis buffer (PBS pH 7.4 (387mM NaCl), 1 mM EDTA, 1X protease inhibitor). Nuclei were lysed via repeated passes through a 23 Gauge needle. The Nuclear lysate was centrifuged at 10,000 x g to remove membrane debris and clarified supernatant was placed atop a 30-50% sucrose cushion and centrifuged at 100,000 x g for 1hr at 4°C. Nucleocapsids were drawn from the 30-50% interface and resuspended in 10 times draw volumes of hypertonic lysis buffer and pelleted via ultracentrifugation at 100,000 x g for 1 hr. The pellet was finally resuspended in 10-20 μL of hypertonic lysis buffer prior to TEM grid preparation and imaging.

To prepare cryoEM samples, 2.5 μL of hypertonic lysis solution containing either purified WT or mutant HCMV nucleocapsids was applied to 200 mesh holey Quantifoil carbon or gold grids, which had been glow discharged for 30 seconds. The sample was flash-frozen in a 50:50 mixture of liquid ethane and propane using a Gatan Vitrobot IV. CryoEM movies were recorded on a K3 direct electron detector in a Titan Krios electron microscope (FEI) equipped with a Gatan Imaging Filter (GIF). The microscope was operated at 300 kV at a nominal magnification of 81,000X and calibrated pixel size of 0.55 Å at the specimen level with the camera in super-resolution mode. Using SerialEM, 2693 and 1261 movies with viral particles were recorded for the WT and mutant capsid samples, respectively. The cumulative electron dose on the sample was ∼45 e^-^/Å^2^.

### 2.7 CryoEM image pre-processing and 3D reconstruction

We performed frame alignment and motion correction to generate a cryoEM micrograph from each movie with cryoSPARC (Punjani et al., 2017). Patch-aligned and dose-weighted micrographs were transferred to Relion 4.0 for further processing and Topaz automated particle picking (Bepler et al., 2019; Scheres, 2012). In total micrographs for the WT sample contained approximately 2966 A-capsids, 1527 B-capsids, and 42 C-capsids, whereas those for the mutant sample contained 1215 A-capsids, 595 B-capsids, and 22 C-capsids. To improve processing speed, particles were extracted and binned eight times prior to 2D image classifications. A 3D Gaussian ball created by back projection of particles without orientation assignments was used as an initial model for 3D classification while imposing icosahedral symmetry. The icosahedral reconstructions of both A- and B-capsids reached a resolution of 17.6 Å.

To improve reconstruction resolution, we carried out sub-particle constructions as described previously (Liu et al., 2019). Briefly, we used Relion’s symmetry expansion method, with I3 symmetry selected, to generate 60 equivalent orientations for each capsid particle with unique Euler angles for rotation, tilt, and psi. Given that there are 5 redundant orientations of each particle about each 5-fold symmetrical vertex, which have identical tilt and psi angles but different in-plane angle, we kept only 1 of each set of 5, to represent each 5-fold vertex sub-particle. These orientations were then used to identify the position of each 5-fold vertex in Cartesian coordinates within the 2D projection images for extraction as new particles totaling 34,836, 17,967, and 482 two-times binned WT A-capsid, B-capsid, and C-capsid and 14,354 and 6,958 for one-time binned pp150-C4 A-capsid and B-capsids, respectively. Further processing was then carried out imposing C5 symmetry onto the newly extracted particles to reach 4.4, 4.4, and 13.6Å for WT A-capsid, B-capsid, and C-capsid and 4.3, and 4.5Å for pp150-C4 mutant A-capsid and B-capsid, respectively.

### 2.8 Ultra-thin sectioning of infected cells and transmission electron microscopy

WT AD169-BAC infected cells were prepared for TEM as described previously (Peng et al., 2010). Briefly, cells were seeded on tissue culture plates for 48 hours and fixed in 2% glutaraldehyde in 1X PBS (pH 7.4). Cells were scraped and thereafter pelleted by low-speed centrifugation. Pellets were subject to 3% uranyl acetate staining following treatment with 1% osmium tetroxide. Samples were dehydrated in ascending ethanol series and embedded in Agar Low Viscosity Resin (Agar Scientific). Samples were sectioned using an UCT ultra microtome (LEICA) with a 35° diamond knife (Diatome, Switzerland). Sections were 55-65 nm thick and collected on 150 mesh hexagonal or slot grids with Formvar-carbon coated support films and stained with saturated uranyl acetate followed by Sato lead citrate on both sides.

To enrich AD169-BAC infected cells for sectioning, fluorescent cells were concentrated via fluorescent activated cell sorting (FACS). Briefly, tissue culture flasks were seeded with mutant AD169-BAC infected cells and grown for 5 days. To prepare for sorting, cells were thoroughly rinsed with PBS (pH 7.4) before trypsinization and incubation at 37° C for several minutes, or until cells detached. Trypsinized cells were resuspended in PBS (pH 7.4) containing 1% bovine serum albumin (BSA) and soybean trypsin inhibitor (ATCC). Cells were pelleted and resuspended in 1% BSA in PBS solution and passed through a 40 um cell strainer (pluriSelect) into FACS culture tubes. Cells were sorted using a FACSAria III High-Speed Cell Sorter (BD Biosciences) gated on GFP. Sorted cells were seeded on two wells of a 6 well plate and supplemented with DMEM growth medium and grown for 2 days prior to fixation.

For nucleocapsid quantitative analysis, TEM imaging was performed using a FEI Titan 80-300 electron microscope operated at 300 kV. Images were recorded on a DE-64 camera (Direct Electron) at a nominal magnification of 10,000 x. Montage micrographs of cell sections were collected using SerialEM (Mastronarde, 2003), by collecting polygons in “view” mode. Defocus was maintained at −40 um during the data acquisition. A, B, and C capsids were annotated on montages using 3dmod in the IMOD package (Kremer et al., 1996). Non-montaged TEM micrographs were recorded on a charged coupled detector (CCD) camera in an FEI Tecnai G2 T20 electron microscope operated at 200 kV to enhance contrast.

## 3. Results

### 3.1 Multipoint mutations at the SCP-pp150 interface produce lethal and attenuating phenotypes

Atomic models of HCMV present a relatively small (∼446 Å^2^) pp150-SCP interface, composed of clusters of complementary hydrophobic or polar amino acids (Fig. 1A-C).(Baxter and Gibson, 2001) From the pp150 face, negatively charged residues (E273, D274, and D276) accommodate a positively charged protrusion from SCP (K46, R49), while aromatic residues from pp150 (F245, Y249) form an Aro-Met-Aro bridge with SCP (M67), and polar uncharged residues from pp150 (N248, N252) contact the C-terminal residues of SCP. We introduced multipoint mutations to substitute each complementary cluster with a non-complementary one (Fig. 1C-D). The impact of each set of mutations on pp150-SCP interaction was first predicted *in silico* using the software package mmCSM-PPI (Rodrigues et al., 2021) (Table S1). Mutations to the F245-M67-Y249 bridge and charged faces were predicted to significantly alter the free energy of association between pp150 and SCP, while those pp150 mutations near the SCP C-terminus (pp150-C4) and negative controls far from the interface (pp150-C5 & C6) were not predicted to impact binding (Table S1).

To validate these structure-based predictions, we transfected ARPE19 cells with AD169-BACmids carrying cluster mutations to either pp150 or SCP and GFP as a positive marker (Fig. 1). As positive control, cells were transfected with wild-type (WT) AD169-BACmids which produced WT virus that spread throughout cell culture as characterized by GFP spread (Fig. 1E-F). In keeping with computational predictions, cluster mutations outside the pp150-SCP interface had no observable impact on viral replication (Fig. 1E), however mutations at any polar or hydrophobic regions of the interface, including near N248 and N252 of pp150, proved lethal to the virus as indicated by lack of GFP spread beyond initially transfected cells 7 days post transfection (Fig. 1E-F). Interestingly, the R240E R251E K255E mutant (pp150-C4) demonstrated fluorescence spread indicative of viral spread albeit with significant attenuation compared to WT HCMV (Fig. 1F). While the lethal mutations were localized to the direct polar or hydrophobic interfaces between pp150 and SCP, R240 and K251 of pp150-C4 only interact with the far CC-terminal region of SCP, while K255 interfaces with MCP, suggesting pp150-C4 is less involved with the SCP interactions. To determine whether it was mutations to the SCP or MCP interface responsible for the attenuation, we generated separate pp150 R240E K251E and pp150 K255E mutants (pp150-C4a and pp150-C4b respectively) to determine whether it was the change to the SCP or MCP adjacent residue(s) leading to the observed phenotype (Fig. 1D-F. Viral replication, as exhibited by GFP spread, showed the R240E K251E mutant to have wild-type kinetics, whereas K255E exhibited attenuated viral spread similar to pp150-C4 (Fig 1F).

### 3.2 Multipoint mutations at the SCP-pp150 interface cause lethal phenotypes and disrupt pp150-capsid interactions

We next sought to determine whether disruption of the pp150-SCP interaction was responsible for the lethal and attenuated phenotypes we observed. To this end, we cloned pp150-hemagglutinin (HA) and FLAG-SCP tagged proteins into a pcDNA3 expression vector. The plasmid DNAs of both WT and mutant genes were co-transfected into ARPE-19 cells and fixed for immunofluorescence assays (IFA) (Fig. 2A). IFAs revealed colocalization of WT SCP to pp150 as cytosolic puncta of transfected ARPE-19 cells (Fig. 2A, yellow). By contrast, plasmids containing lethal SCP or pp150 mutants exhibited delocalized signal dispersed throughout the cytosol (Fig. 2A). The pp150-C4 mutant virus demonstrated SCP colocalization similar to WT, indicating pp150-C4 readily associates with SCP despite the attenuated replication. This finding is consistent with the notion that the K255E mutation drives the attenuation via disruption of the pp150-MCP interface. To corroborate the IFA results, the plasmids were co-transfected into HEK-293 cells for co-immunoprecipitation assays. In WT controls, FLAG-tagged SCP was readily detected in the immunoblot; but the lethal mutations showed no FLAG signal (Fig. 2B). By comparison, pp150-C4 demonstrated SCP signal consistent with WT, further supporting our hypothesis that the interaction is not disrupted (Fig. 2B). These findings together demonstrate that lethal cluster mutations are those that disrupt the pp150-SCP interface and that disruptions to other interfaces lead to attenuating mutations.

**Figure 2.**
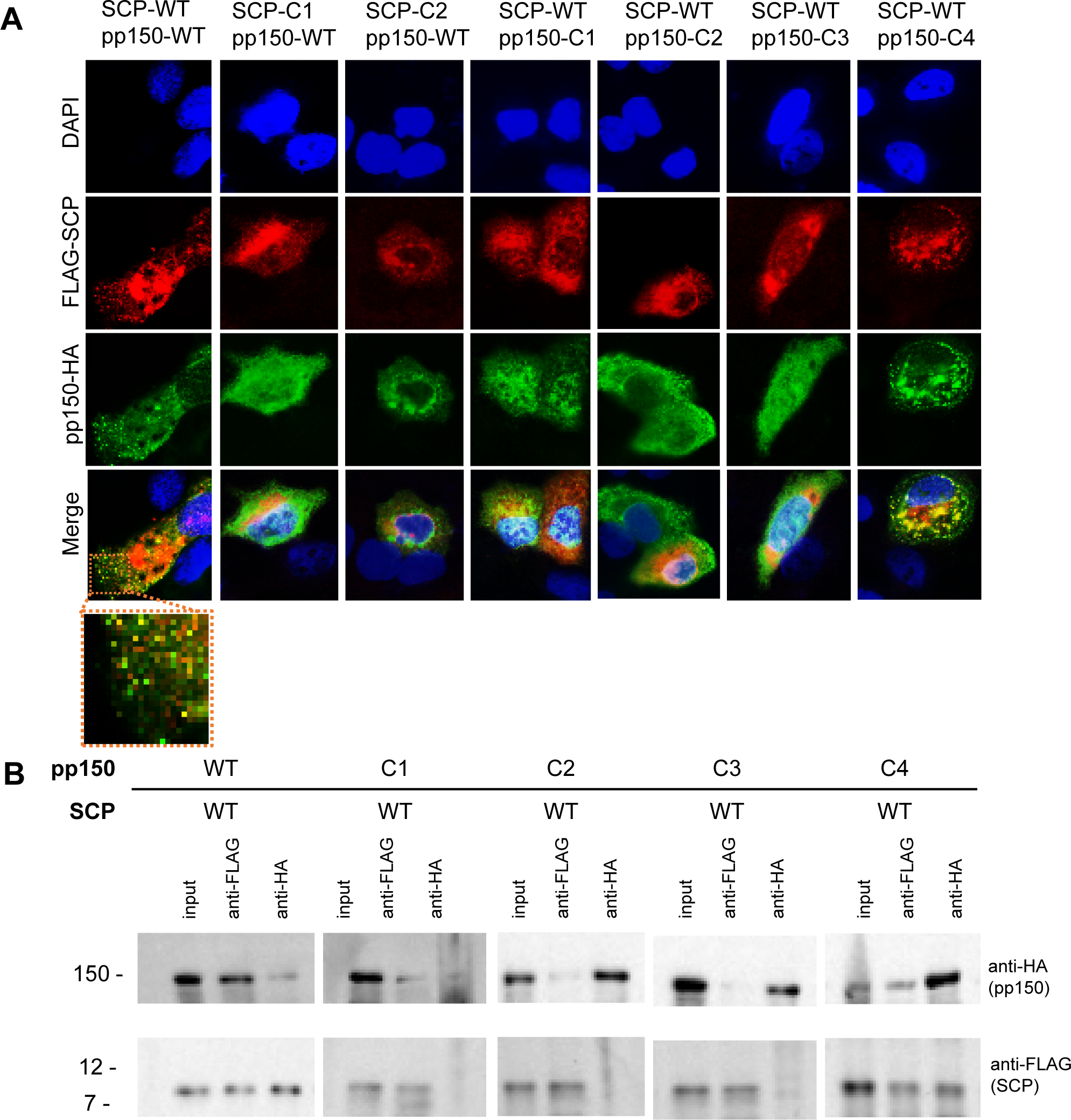
Characterization of mutant pp150 and SCP interactions. (A) Immunofluorescence assays of ARPE-19 cells transfected with either WT or cluster mutations on pp150 or SCP showing colocalization between anti-HA (green) and anti-FLAG (red) with DAPI stained nuclei (blue). (B) Coimmunoprecipitation (CoIP) assays showing interactions of pp150 cluster mutants with WT SCP. Protein extracts were precipitated with anti-HA or anti-FLAG antibodies and resolved by SDS-PAGE. Protein-protein interactions were immunodetected using anti-HA and anti-FLAG antibodies.

### 3.3 Pp150-attenuated mutants maintain genome packaging and viral maturation

To characterize the intracellular consequences of the pp150-C4 mutant’s impaired pp150-MCP interaction on viral assembly and replication, we enriched pp150-C4 mutant HCMV infected cells using fluorescence activated cell sorting (FACS) gated on GFP signal. These pp150-C4 infected cells were fixed, embedded in resin, and used to generate ultra-thin sections (∼55 nm) for TEM analysis. These ultra-thin sections were transferred to formvar-carbon coated TEM grids, stained for large-field of view transmission electron microscopy with Montaging in SerialEM (Mastronarde, 2005). Nuclear membranes were identified and nucleocapsids populations were categorized into three groups including (A) empty, (B) scaffold protein-containing, and (C) genome-filled capsids, based on appearance when observed by electron microscopy (Fig. 4A). WT nuclei appeared with greater proportions of B and C capsids compared to A and this distribution was similar in pp150-C4 infected cells (Fig. 3A-B). In addition to having nucleocapsid distribution similar to WT, pp150-C4 infected cells had vAC associated viral products including enveloped C-capsids, suggesting efficient viral maturation (Fig. 3C).

**Figure 3.**
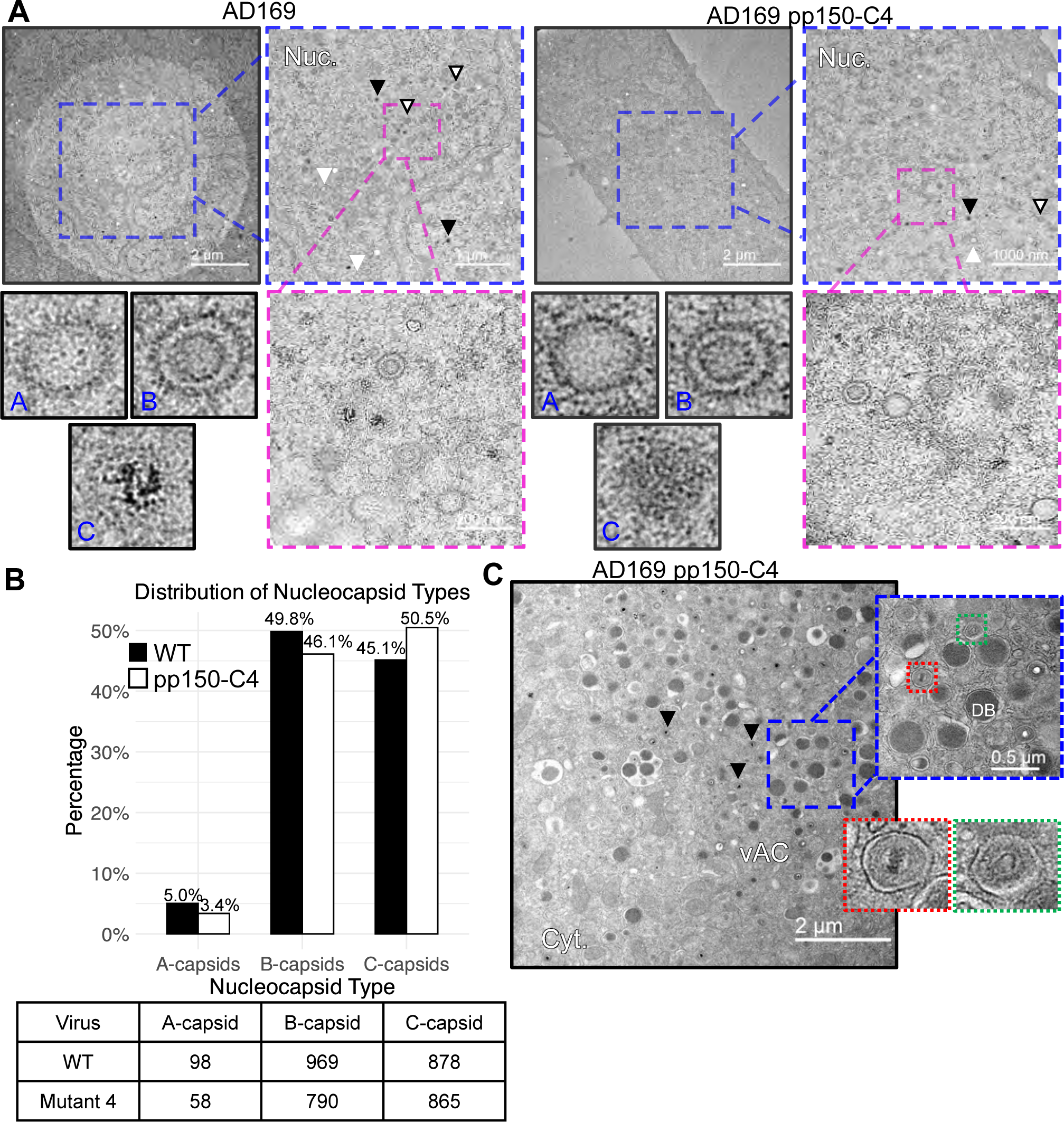
Thin section TEM of ARPE-19 cells infected with WT and mutant HCMV. (A) Representative TEM micrographs of WT (left) and AD169 pp150-C4 infected cells (right) with zoomed in views highlighting different capsid types, with some A-(white arrowheads), B-(white with black outline arrowheads), and C-capsids (black arrowheads) indicated. (B) Distribution of A-, B-, and C-nucleocapsids as percentages of total counted and totals in accompanying table (right). (D) Representative micrograph of vAC from AD169 pp150-C4 infected cells with zoomed in views of particles undergoing envelopment (right, top) along with dense bodies (DB) and examples of enveloped particles (right, bottom).

**Figure 4.**
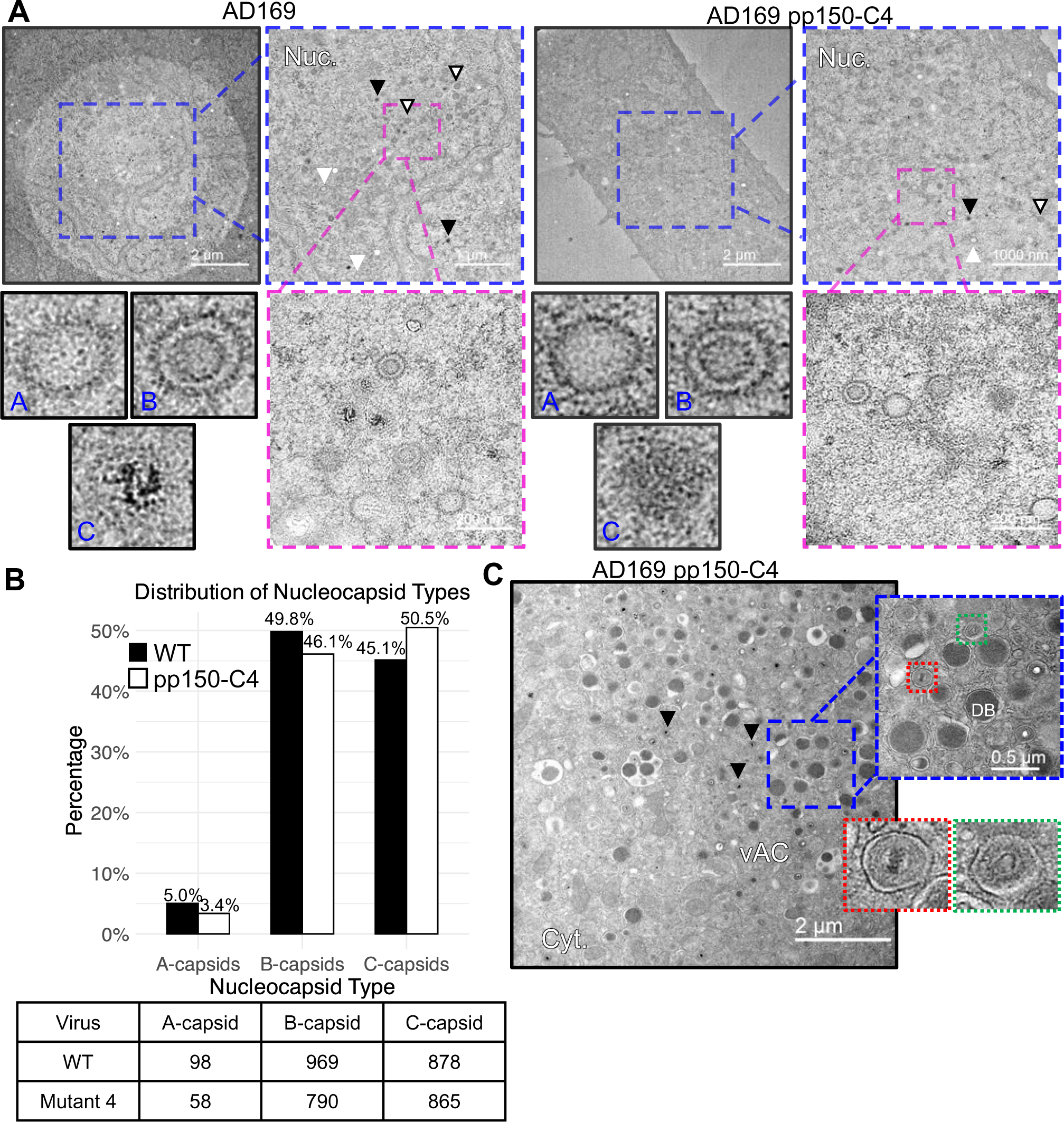
3D reconstructions of nucleocapsids reveal variable tegumentation. (A) Representative cryoEM images of A-, B-, and C-capsids purified from WT AD169 (left) and AD169 pp150-C4 mutant infected cells (right). White arrows point to empty space between MCP towers. (B) Reconstruction of WT (left) and AD169 pp150-C4 A- and B-capsids colored by distance from center of capsid. (C) Vertex sub-particle reconstruction of particles from **B**, colored by capsid proteins as indicated to convey nucleocapsid composition, viewed from top (top) and side (bottom). (D) Low resolution vertex sub-particle reconstruction from the WT C-capsids showing lacking pp150 density. (E) Representative cryoEM images demonstrating fuzzy tegumentation (black arrow) around B-(top) and A-capsids (bottom) with percentages from total B- and A-capsids observed with tegument and (F) a vertex sub-particle generated from such particles (right) showing the presence of pp150 density (cyan) and asymmetric unit with distribution of pp150 and tRNA (right). (G) Representative images of pp150-C4 mutant capsids with fuzzy tegumentation (black arrows).

### 3.4 pp150 is translocated across the nuclear membranes to bind capsids in the nucleus

Due to lack of recognizable nuclear localization signal on the pp150 protein sequence and lack of characterized chaperones, it is unclear how pp150 is translocated into the nucleus and whether it binds to the capsids there or in the cytosol where it also accumulates (Mitra et al., 2021; Tandon and Mocarski, 2008). Indeed previous studies have demonstrated the nucleocapsids capacity for nuclear egress without pp150 (Tandon and Mocarski, 2008). After finding pp150-C4-HCMV had no obvious defects in vAC formation, we sought to determine whether pp150 was being properly incorporated into the nucleocapsids of the mutant virus. The low replication efficiency of pp150-C4 virus complicated efforts to purify complete virions from culture media and necessitated a modified culture procedure to purify nucleocapsids from infected cells. Similar to WT HCMV (Fig. 4A-C, left), the pp150-C4 HCMV nucleocapsids purified from cell nuclei were predominantly A-capsid or B-capsids (Fig 4A-C, right), consistent with previous findings (Li et al., 2023; Tandon et al., 2015)(Li et al., 2023; Tandon et al., 2015).

Initial cryoEM capsid reconstructions from WT and pp150-C4 mutant samples showed no obvious densities at locations occupied by pp150 in WT virions (Fig. 4B-C). 3D reconstruction from a small subset of genome containing C-capsids also lacked pp150 densities (Fig. 4E). While these findings were consistent with previously reported structures (Li et al., 2023), closer inspection of cryoEM images of nucleocapsid revealed conspicuously empty spaces between MCP turrets, which differed from cryoEM micrographs of virions (Fig. 4A). Using cryoEM images from virions as a template, we identified a visually distinguishable subset of predominantly A-capsids and some B-capsids with density corresponding to the tegument (Fig. 4E). When subjected to in-depth data processing, this subset of images yielded a 3D reconstruction with cryoEM densities corresponding to pp150 and tRNAs similar to those observed in previously published structures (Fig. 4E) (Liu et al., 2021). While insufficient numbers of tegument bound particles were present in the pp150-C4 dataset for 3D reconstruction to resolve similar densities, capsid images with tegumentation like that of the WT were observed, suggesting pp150-capsid association in the mutant (Fig. 4F).

## 4. Discussion

Amongst herpesviruses, virion assembly is a tightly regulated process driven by virus-specific proteins, making it an attractive target for drug development. The β-herpesvirus specific tegument protein pp150 is integral to cytoplasmic assembly and requires nucleocapsid incorporation to carry out its functions (Mitra et al., 2021; Tandon and Mocarski, 2008; Yu et al., 2017)(Mitra et al., 2021; Tandon and Mocarski, 2008; Yu et al., 2017). Recently, the detailed interactions between pp150 and the nucleocapsid were elucidated with the publication of several high-resolution cryoEM structures of HCMV virions (Li et al., 2021; Yu et al., 2017). The current study leveraged these structures to carry out the mutational analysis of several well-conserved domains essential to the pp150-SCP interaction, identify a mutation which considerably attenuates viral replication, and confirm that pp150 associates with viral capsids within the nucleus of infected cells.

Previous functional investigations of pp150 have largely focused on CR1, CR2, and the highly conserved cysteine tetrad which were recognized decades ago (Baxter and Gibson, 2001; Mitra et al., 2021). Indeed, peptide mimics of CR2 have shown promise as inhibitors of pp150-capsid association, however the mechanism of inhibition remains unclear as CR1 and CR2 do not obviously interface with capsid proteins according toto available CMV structures (Mitra et al., 2021). We found that substitutions in the conserved polar or hydrophobic domains directly involved in the pp150-SCP interface were detrimental to viral replication and these effects could be directly attributed to disruption of pp150-SCP association (Figs. 1 & 2). No mutations presented here overlapped with CR1 or CR2, however, many of the residues selected were well conserved across SCMV and MCMV (Fig. 1B). Thus, we suspect these conserved residues are essential for capsid association of pp150 and its homologues across cytomegaloviruses of various species, but this will require further investigation.

Our study also hints that pp150’s interaction with SCP begins earlier than previously thought. IFA and coimmunoprecipitation assays revealed pp150 binds cytoplasmic SCP in cells co-transfected with both proteins (Fig. 2A). Considering pp150 and SCP are expressed at similar timepoints during viral infection (Ye et al., 2020), it is conceivable that these interactions occur in virus infected cells as well. Without an obvious nuclear translocation signal or chaperone, it is unclear how pp150 reaches the nucleus. As SCP is thought to be shuttled to the nucleus by scaffolding proteins along with MCP (Gibson, 2008), pp150’s cytoplasmic association to this complex would provide a plausible mechanism for nuclear transport, however this hypothesis will require future investigation.

Despite evidence suggesting pp150 tegumentation occurs in the nuclei of infected cells, previous structural investigations of genome-free, nucleus derived HCMV nucleocapsids indicated the absence of pp150 (Li et al., 2023; Trus et al., 1999). We likewise observed that most nucleus derived capsids in our dataset lacked tegumentation in cryoEM images and 3D reconstructions (Fig. 4). Curiously, C-capsids purified from the nucleus, typically considered the most likely candidates for pp150 association, also lacked pp150 tegumentation as revealed by cryoEM reconstruction. Approximately 8% and 4% of A- and B-capsids respectively had tegumentation, consisting of pp150 and tRNA, observable in cryoEM images (Fig. 4E). As nucleus derived C-capsids are notoriously unstable, and in infected cells, A-capsids compose only a small percentage of viral particles (Fig. 3 and Tandon et al., 2015), one may reasonably assume that many A-capsids in our purified sample arose from C-capsids whose genomes prematurely ejected. Therefore, the greater proportion of tegumented A-capsids we observed may reflect preferential association of pp150 to C-capsids over B-capsids.

Prior studies demonstrated that pp150-deletion renders HCMVHCMV unable to mature beyond the cytoplasmic stage, with few cytoplasmic capsids being observed and a highly vesiculated vAC (Tandon and Mocarski, 2008). By contrast, our pp150-C4 mutant HCMV appeared capable of efficient genome packaging, nuclear egress and vAC formation despite significant attenuation to viral spread (Figs. 1 & 3). This would appear to suggest that the effects of such a mutation are either too subtle for detection in our TEM surveys or that pp150-C4 mutants are impacted at another point during their replication cycle. The large number of complex processes comprising capsid maturation and incomplete characterization of pp150’s role throughout makes it difficult to pinpoint where along that process this mutation impacts. Nonetheless, the work reported here represents the first time structure-guided design and BAC mutagenesis has led to identification of lethal, as well as an attenuated HCMV mutant and raises important questions regarding the role of the pp150-SCP and pp150-MCP interactions in viral replication. The latter produces far less infectious virus compared the wild-type virus and warrants further study in the pursuit as candidate of live attenuated vaccine.

## 5. Conclusion

In summary, guided by atomic structure of HCMV, we demonstrated the sensitivity of the pp150-SCP interface to mutations, while also providing the first structural evidence of pp150 binding to capsids inside nucleus. While many of these interacting residues are essential to HCMV replication, we identified a point mutation that significantly attenuates intercellular transmission of the virus in vitro, which opens the door for further exploration as to develop a live attenuated HCMV vaccine.

## CRediT authorship contribution statement

**Alex Stevens**: Overall experimental design and execution of the electron microscopy-related components, data analysis and result interpretation, manuscript drafting and figure preparation, paper editing. **Ruth Cruz-cosme**: BAC-related experiments, co-IP, and western blot assays. **Najealicka Armstrong**: molecular cloning to generate plasmids and immunofluorescence assays. **Qiyi Tang**: Design and supervision of BAC-related experiments, co-IP experiments, fluorescence microscopy, and manuscript editing. **Z. Hong Zhou**: Overall project design and supervision of execution, research fund acquisition and management, manuscript draft and editing.

## Declaration of competition interest

The authors declare that they have no known competing financial interests or personal relationships that could have appeared to influence the work reported in this paper.

## Data availability

The cryoEM structures reported here are deposited at the Electron Imaging Database under the accession code of EMD-XXXXX (mutant A-capsid), EMD-XXXXX (mutant B-capsid), EMD-XXXXX (WT A-capsid), EMD-XXXXX (WT B-capsid), and EMD-XXXXX (WT C-capsid).

## Acknowledgements

We thank Sergey Ryazantsev & Chunni Zhu for the cell embedding and thin-sectioning service. This work was supported by grants from the US National Institutes of Health (R01DE028583 to ZHZ). A.S. Received support from NIH Ruth L. Kirschstein National Research Service Award AI007323. We acknowledge the use of resources in the Electron Imaging Center for NanoSystems (EICN) supported by UCLA, National Science Foundation (DMR-1548924 and DBI-1338135) and National Institutes of Health (S10RR23057 and U24GM116792). We acknowledge support from the UCLA AIDS Institute, the UCLA Brain Research Institute Electron Microscopic Core, the James B. Pendleton Charitable Trust and the McCarthy Family Foundation.

---

**Table S1.** *In-silico* mutational analysis of pp150-SCP interface. Table from Figure 1D extended to include predicted change to binding affinity (ΔΔG) in kcal/mol by either SCP or pp150 mutant clusters. Decreased affinity values are negative and colored red and increased affinity are positive colored in blue.

| Mutant | Mutations | Interface(s) | Phenotype | mmCSM-PPI<br>$\Delta\Delta G$ (kcal/mol) |
| --- | --- | --- | --- | --- |
| WT | N/A | N/A | Healthy | NA |
| SCP-C1 | M67W | pp150 | Lethal | -0.91 |
| SCP-C2 | K46E, R49E | pp150 | Lethal | -2.09 |
| pp150-C1 | E273K, D274K, D276K | SCP | Lethal | +0.09 |
| pp150-C2 | N248E, N252E | SCP | Lethal | -0.49 |
| pp150-C3 | F245A, Y249A | SCP | Lethal | -1.08 |
| pp150-C4 | R240E, R251E, K255E | SCP, MCP | Attenuated | -0.03 |
| pp150-C4a | R240E, R251E | SCP | Healthy | 0.05 |
| pp150-C4b | K255E | MCP | Attenuated | 0 |
| pp150-C5 | K233D, K237D | N/A | healthy | -0.03 |
| pp150-C6 | D262K, D265K | N/A | healthy | -0.03 |

## References

AuCoin, D.P., Smith, G.B., Meiering, C.D., Mocarski, E.S., 2006. Betaherpesvirus-Conserved Cytomegalovirus Tegument Protein ppUL32 (pp150) Controls Cytoplasmic Events during Virion Maturation, Journal of Virology, pp. 8199–8210.

Bauer, D.W., Huffman, J.B., Homa, F.L., Evilevitch, A., 2013. Herpes Virus Genome, The Pressure Is On, Journal of the American Chemical Society, pp. 11216–11221.

Baxter, M.K., Gibson, W., 2001. Cytomegalovirus Basic Phosphoprotein (pUL32) Binds to Capsids In Vitro through Its Amino One-Third, Journal of Virology, pp. 6865–6873.

Bepler, T., Morin, A., Rapp, M., Brasch, J., Shapiro, L., Noble, A.J., Berger, B., 2019. Positive-unlabeled convolutional neural networks for particle picking in cryo-electron micrographs, Nature Methods. Springer US, pp. 1153–1160.

Borst, E.M., Harmening, S., Sanders, S., Caragliano, E., Wagner, K., Lenac Roviš, T., Jonjić, S., Bosse, J.B., Messerle, M., 2022. A Unique Role of the Human Cytomegalovirus Small Capsid Protein in Capsid Assembly, in: Damania, B. (Ed.), mBio. American Society for Microbiology.

Borst, E.M., Wagner, K., Binz, A., Sodeik, B., Messerle, M., 2008. The Essential Human Cytomegalovirus Gene UL52 Is Required for Cleavage-Packaging of the Viral Genome, Journal of Virology, pp. 2065–2078.

Brandariz-Nuñez, A., Liu, T., Du, T., Evilevitch, A., 2019. Pressure-driven release of viral genome into a host nucleus is a mechanism leading to herpes infection, eLife, pp. 1–20.

Close, W.L., Anderson, A.N., Pellett, P.E., 2018. Betaherpesvirus Virion Assembly and Egress, pp. 167–207.

Dai, X., Gong, D., Xiao, Y., Wu, T.-T., Sun, R., Zhou, Z.H., 2015. CryoEM and mutagenesis reveal that the smallest capsid protein cements and stabilizes Kaposi’s sarcoma-associated herpesvirus capsid, Proceedings of the National Academy of Sciences, pp. E649–E656.

Dai, X., Yu, X., Gong, H., Jiang, X., Abenes, G., Liu, H., Shivakoti, S., Britt, W.J., Zhu, H., Liu, F., Zhou, Z.H., 2013. The Smallest Capsid Protein Mediates Binding of the Essential Tegument Protein pp150 to Stabilize DNA-Containing Capsids in Human Cytomegalovirus, in: Gibson, W. (Ed.), PLoS Pathogens, p. e1003525.

Dai, X., Zhou, Z., 2014. Purification of Herpesvirus Virions and Capsids, Bio-Protocol, pp. 2–6.

Dai, X., Zhou, Z.H., 2018. Structure of the herpes simplex virus 1 capsid with associated tegument protein complexes, Science, p. eaao7298.

Gibson, W., 2008. Structure and Formation of the Cytomegalovirus Virion, in: Shenk, T.E., Stinski, M.F. (Eds.), Current Topics in Microbiology and Immunology. Springer, Berlin, pp. 187–204.

Kremer, J.R., Mastronarde, D.N., McIntosh, J.R., 1996. Computer Visualization of Three-Dimensional Image Data Using IMOD, Journal of Structural Biology, pp. 71–76.

Li, Z., Pang, J., Dong, L., Yu, X., 2021. Structural basis for genome packaging, retention, and ejection in human cytomegalovirus, Nature Communications. Springer US, p. 4538.

Li, Z., Pang, J., Gao, R., Wang, Q., Zhang, M., Yu, X., 2023. Cryo-electron microscopy structures of capsids and in situ portals of DNA-devoid capsids of human cytomegalovirus, Nature Communications. Springer US, p. 2025.

Liu, Y.-t., Strugatsky, D., Liu, W., Zhou, Z.H., 2021. Structure of human cytomegalovirus virion reveals host tRNA binding to capsid-associated tegument protein pp150. Nature Communications, 1-9.

Liu, Y.-T.T., Jih, J., Dai, X., Bi, G.-Q.Q., Zhou, Z.H., 2019. Cryo-EM structures of herpes simplex virus type 1 portal vertex and packaged genome, Nature. Springer US, pp. 257–261.

Manicklal, S., Emery, V.C., Lazzarotto, T., Boppana, S.B., Gupta, R.K., 2013. The “Silent” Global Burden of Congenital Cytomegalovirus, Clinical Microbiology Reviews, pp. 86–102.

Mastronarde, D.N., 2003. SerialEM: A program for automated tilt series acquisition on Tecnai microscopes using prediction of specimen position, Microscopy and Microanalysis, pp. 1182–1183.

Mastronarde, D.N., 2005. Automated electron microscope tomography using robust prediction of specimen movements. J Struct Biol 152, 36–51.

Mitra, D., Hasan, M.H., Bates, J.T., Bidwell, G.L., Tandon, R., 2021. Tegument Protein pp150 Sequence-Specific Peptide Blocks Cytomegalovirus Infection, Viruses. MDPI, p. 2277.

Muller, C., Alain, S., Baumert, T.F., Ligat, G., Hantz, S., 2021. Structures and Divergent Mechanisms in Capsid Maturation and Stabilization Following Genome Packaging of Human Cytomegalovirus and Herpesviruses, Life, p. 150.

Peng, L., Ryazantsev, S., Sun, R., Zhou, Z.H., 2010. Three-Dimensional Visualization of Gammaherpesvirus Life Cycle in Host Cells by Electron Tomography, Structure, pp. 47–58.

Punjani, A., Rubinstein, J.L., Fleet, D.J., Brubaker, M.A., 2017. cryoSPARC: algorithms for rapid unsupervised cryo-EM structure determination, Nature Methods, pp. 290–296.

Rodrigues, C.H.M., Pires, D.E.V., Ascher, D.B., 2021. mmCSM-PPI: predicting the effects of multiple point mutations on protein–protein interactions, Nucleic Acids Research. Oxford University Press, pp. W417–W424.

Sae-Ueng, U., Liu, T., Catalano, C.E., Huffman, J.B., Homa, F.L., Evilevitch, A., 2014. Major capsid reinforcement by a minor protein in herpesviruses and phage. Nucleic Acids Research 42, 9096–9107.

Scheres, S.H.W., 2012. RELION: Implementation of a Bayesian approach to cryo-EM structure determination, Journal of Structural Biology. Elsevier Inc., pp. 519–530.

Tandon, R., Mocarski, E., Conway, J., 2015. The A, B, Cs of Herpesvirus Capsids, Viruses, pp. 899–914.

Tandon, R., Mocarski, E.S., 2008. Control of Cytoplasmic Maturation Events by Cytomegalovirus Tegument Protein pp150, Journal of Virology, pp. 9433–9444.

Tandon, R., Mocarski, E.S., 2011. Cytomegalovirus pUL96 Is Critical for the Stability of pp150-Associated Nucleocapsids, Journal of Virology, pp. 7129–7141.

Trus, B.L., Gibson, W., Cheng, N., Steven, A.C., 1999. Capsid Structure of Simian Cytomegalovirus from Cryoelectron Microscopy: Evidence for Tegument Attachment Sites, Journal of Virology, pp. 4530–4530.

Wang, D., Shenk, T., 2005. Human cytomegalovirus UL131 open reading frame is required for epithelial cell tropism. J Virol 79, 10330–10338.

Wang, J., Yuan, S., Zhu, D., Tang, H., Wang, N., Chen, W., Gao, Q., Li, Y., Wang, J., Liu, H., Zhang, X., Rao, Z., Wang, X., 2018. Structure of the herpes simplex virus type 2 C-capsid with capsid-vertex-specific component, Nature Communications. Springer US, p. 3668.

Warden, C., Tang, Q., Zhu, H., 2011. Herpesvirus BACs: Past, Present, and Future, Journal of Biomedicine and Biotechnology, pp. 1–16.

Ye, L., Qian, Y., Yu, W., Guo, G., Wang, H., Xue, X., 2020. Functional Profile of Human Cytomegalovirus Genes and Their Associated Diseases: A Review. Frontiers in Microbiology 11, 1–13.

Yu, X., Jih, J., Jiang, J., Zhou, Z.H., 2017. Atomic structure of the human cytomegalovirus capsid with its securing tegument layer of pp150, Science, p. eaam6892.

Zuhair, M., Smit, G.S.A., Wallis, G., Jabbar, F., Smith, C., Devleesschauwer, B., Griffiths, P., 2019. Estimation of the worldwide seroprevalence of cytomegalovirus: A systematic review and meta-analysis, Reviews in Medical Virology, p. e2034.

